# Delegating sex: differential gene expression in stolonizing syllids uncovers the hormonal control of reproduction in Annelida

**DOI:** 10.1101/271783

**Authors:** Patricia Álvarez-Campos, Nathan J. Kenny, Aida Verdes, Rosa Fernández, Marta Novo, Gonzalo Giribet, Ana Riesgo

**Author notes:** Present address: Stem Cells, Development and Evolution, Institut Jacques Monod, CNRS UMR 7592, Université Paris-Diderot;15 rue Hélène Brion 75205 Paris cedex 13, France. Senior authors (Gonzalo Giribet, Ana Riesgo).

## Abstract

Stolonization in syllid annelids is a unique mode of reproduction among animals. During the breeding season, a structure resembling the adult but containing only gametes, called stolon, is formed at the posterior end of the animal. When the stolons mature, they detach from the adult and the gametes are released into the water column. The process is synchronized within each species, and it has been reported to be under environmental and endogenous control, probably via endocrine regulation. To further understand the reproduction in syllids and to elucidate the molecular toolkit underlying stolonization, we generated Illumina RNA-seq data from different tissues of reproductive and non-reproductive individuals of *Syllis magdalena*, and characterized gene expression during the stolonization process. Several genes involved in gametogenesis (*ovochymase*, *vitellogenin*, *testis-specific serine/threonine-kinase*), immune response (*complement receptor 2*), neuronal development (*tyrosine-protein kinase Src42A*), cell proliferation (*alpha-1D adrenergic receptor*), and steroid metabolism (*hydroxysteroid dehydrogenase* 2) were found differentially expressed in the different tissues and conditions analyzed. In addition, our findings suggest that several neurohormones, such as methyl farnesoate, dopamine and serotonin, might trigger the stolon formation, the correct maturation of gametes and the detachment of stolons when gametogenesis is complete. The process seems to be under circadian control, as indicated by the expression patterns of *r-opsins*. Overall, our results shed light into the genes that orchestrate the onset of gamete formation, and improve our understanding of how some hormones, previously reported to be involved in reproduction and metamorphosis processes in other invertebrates, seem to also regulate reproduction via stolonization.

## Introduction

Annelids in the family Syllidae have a remarkable reproductive strategy, which has attracted the attention of many biologists (e.g. Nygren 1999 and references herein). Syllids exhibit epitoky, which largely implies morphological changes associated with reproduction (Malaquin 1893), and can be further divided into a variety of reproductive modes. In all epitokous modes, there are two states: the sexually immature worm, called an “atoke”, and the sexually mature worm, or “epitoke”. Among the epitokous types of reproduction, one of the most common is epigamy, which is not exclusive to syllids, where the entire atoke transforms into the epitoke, developing swimming chaetae, enlarging its eyes and undergoing changes in musculature (Wissocq 1970; Daly 1975; Garwood 1991). One of the most extreme types of epitokous reproduction is squizogamy or stolonization, where only a part of the individual transforms into an epitokal sexual stage, either by generating new segments or by differentiating pre-existing ones (Franke 1999). When the breeding season approaches, the syllid atoke (or stock) starts to develop a peculiar structure at the end of its body, that resembles the adult and is known as the stolon (Agassiz 1863) (Fig. 1). The stolons possess several features similar to the stock, such eyes and antennae, but are filled with gametes (Figs. 1 and 2A–E), as their brief existence is exclusively devoted to mating, followed by death (Franke 1999). The stock produces and transfers the gametes to the stolon, which is released from the stock when mature (with developed eyes and antennae) (Figs. 1 and 2E), and swims to the surface to spawn (Potts 1911; Mesnil and Caullery 1919). The pelagic stolon releases gametes into the water column, via the nephridiopores in the case of sperm, and through rupture of the body wall for the eggs (Franke 1980; Wissocq 1966, 1970; Schroeder and Hermans 1975; Okada 1937; Durchon 1951, 1952, 1959). Finally, before or after stolon detachment (depending on the species), the stock regenerates the lost final segments (e.g., Marion and Bobretsky 1875; Michel 1898; Okada 1929) (Fig. 1 and 2F).

**Figure 1.**
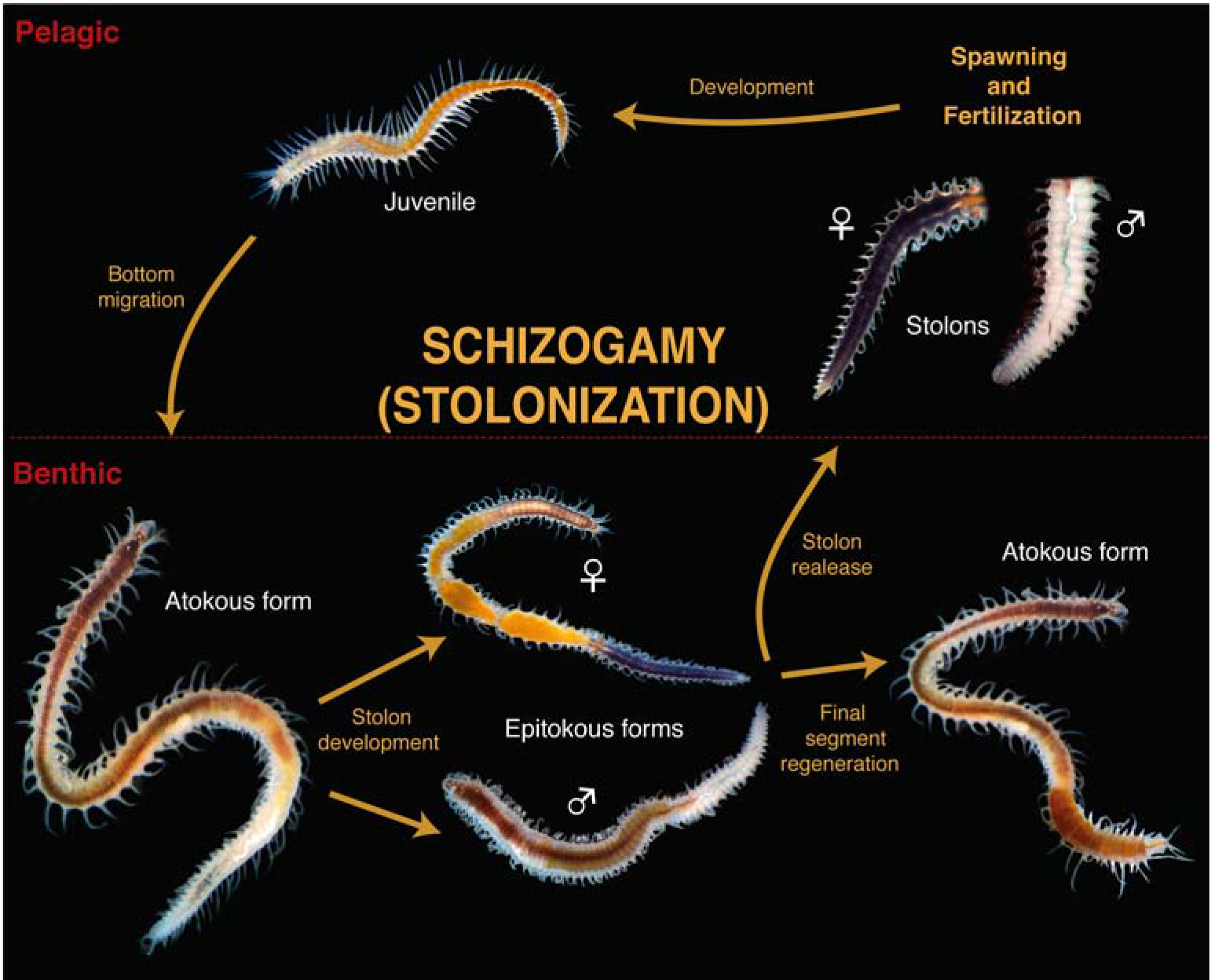
Syllinae schizogamous reproductive cycle (stolonization) using light microscope pictures of *Syllis magdalena*.

The molecular toolkit involved in annelid reproduction is still far from being understood, although studies in several annelid species have shed some light into the matter. For instance, genes involved in pheromone production that are essential for mate recognition and spawning, such as *Temptin* and *Attractin*, and those involved in gametogenesis or fertilization, such as *Fertilin* or *Acrosin*, have been identified in *Spirobranchus* (*Pomatoceros*) *lamarckii, Hormogaster samnitica* and *H. elisae* (e.g., Kang et al. 2002; Rivera et al. 2005; Takahashi et al. 2009; Novo et al. 2013). It is also well-known that the germline specification in the marine annelids *Alitta virens, Platynereis dumerilii, Capitella teleta* or *Hermodice carunculata* involves the expression of several genes including *vasa*, *nanos*, and *piwi* during embryogenesis, and that *vitellogenin* (*Vtg*) is required for yolk formation in the oocyte (Hafer et al.1992; Dill and Seaver 2008; Rebscher et al. 2007; Thamm and Seaver 2008; Giani et al. 2011; Mehr et al. 2015; Schenk et al. 2016). Interestingly, a recent study has reported the potential involvement of the sesquiterpenoid methyl farnesoate (MF), the brain neurohormone that directly regulates *Vtg* in yolk production of *P. dumerilii* females, therefore influencing the correct development of oocytes (Schenk et al. 2016). Particularly, a decrease in MF levels in the brain of *P. dumerilii* during reproduction allowed oocyte maturation but suppressed normal somatic functions and caudal regenerative capacities (Schenk et al. 2016). In crustaceans, MF has been showed to play essential roles in development and reproduction (Xie et al. 2016), similar to the role of Juvenile Hormone (JH) in insects (Riddiford 1994; Wyatt and Davey 1996). Other hormones have also been proposed to play essential roles in annelids reproduction, such as the prostomium (i.e. first pre-oral segment of the animal) hormone 8,11,14-eicosatrienoic acid, which seems to be responsible for sperm maturation and spawning in *Arenicola marina* males (Bentley 1985; Bentley et al. 1990; Pace and Bentley 1992).

Similarly, it has been proposed that the stolonization process in syllids is under hormonal control, following endogenous circadian and circalunar rhythms influenced by exogenous factors, including annual photoperiod, temperature or moon cycles (Franke 1986a; 1999). It has been hypothesized that during the summer time, with long days and high temperatures, a stolonization-promoting hormone produced in the prostomium is secreted to control a second stolonization-suppressing hormone produced in the proventricle (i.e. specialized structure of the digestive tract), allowing the initiation of stolonization (Franke 1999). In contrast, during winter, when days are short and temperatures low at high latitudes, the proventricle is not controlled by the prostomium, and the proventricular stolonization-suppressing hormone then inhibits stolonization (e.g., Abeloos 1950; Durchon 1952, 1959; Durchon and Wissocq 1964; Franke 1980, 1981, 1983a, b, 1985, 1999; Heacox 1980; Heacox and Schroeder 1982; Franke and Pfannenstiel 1984; Verger-Bocquet 1984). Hormonal factors have also been suggested to drive the sexual differentiation of the stolon (Franke 1980; Heacox and Schroeder 1982), in particular the female stolon, given that it seems that male stolon differentiation occurs autonomously, whereas female stolon differentiation may depend on hormone release by male stolons (Franke 1999). However, no candidate hormone has been proposed to control reproduction and regeneration processes in syllids, although it seems clear that there might be several involved, not only in the brain, but also in the proventricle (e.g., Schroeder and Hermans 1975; Franke 1999; Weidhase et al. 2016)

In summary, although molecular mechanisms underlying reproduction are relatively well studied in a few annelids (e.g., Kang et al. 2002; Thamm and Seaver 2008; Giani et al. 2011; Novo et al. 2013; Schenk et al. 2016), the molecular toolkit involved in the stolonization process of syllids has not been examined yet. Thus, our aim in the present study is to provide a first glimpse into the gene expression patterns occurring during the stolonization process in the syllid species *Syllis magdalena*. To achieve this goal, we have pursued four main objectives: 1) to characterize molecularly and morphologically the stolonization process in the target species; 2) to provide a detailed description of the genes potentially involved in the triggering of stolonization and the formation/releasing of stolons and gametes, through differential gene expression analyses of reproductive and non-reproductive individuals in different tissues; 3) to understand the evolution of selected candidate genes with major roles in the reproductive processes of the phylum Annelida; and 4) to investigate if the molecular signal that determines when to divert resources from somatic functions to reproduction is the same across annelids (i.e., synthesis of methyl farnesoate).

## Results and Discussion

### General morphology and ultrastructure of the stolons in Syllis magdalena

The stolons of *S. magdalena* were dicerous, with two pairs of red eyes and one pair of antennae formed at the beginning of the stolonization process (Fig. 2A–E, 3A–B), similar to the process observed in *Syllis amica* (see Wissoq 1970) but different to the late formation of head structures in *Syllis gracilis* (see Pettibone 1963) or *Syllis hyalina* (see Malaquin 1893). Natatorial capillary chaetae were not developed during the stages in which the stolon was attached to the stock. Before stolon detachment, the stock completely regenerates the final part of the body that was transformed during the stolon formation (Fig. 2F). Female stolons were purple, completely full of oocytes arranged around the through-gut (Fig. 2A, C, E, 3A–B). Male stolons were white, completely full of spermatogonia, and also arranged around the gut (Fig. 2B, D).

**Figure 2.**
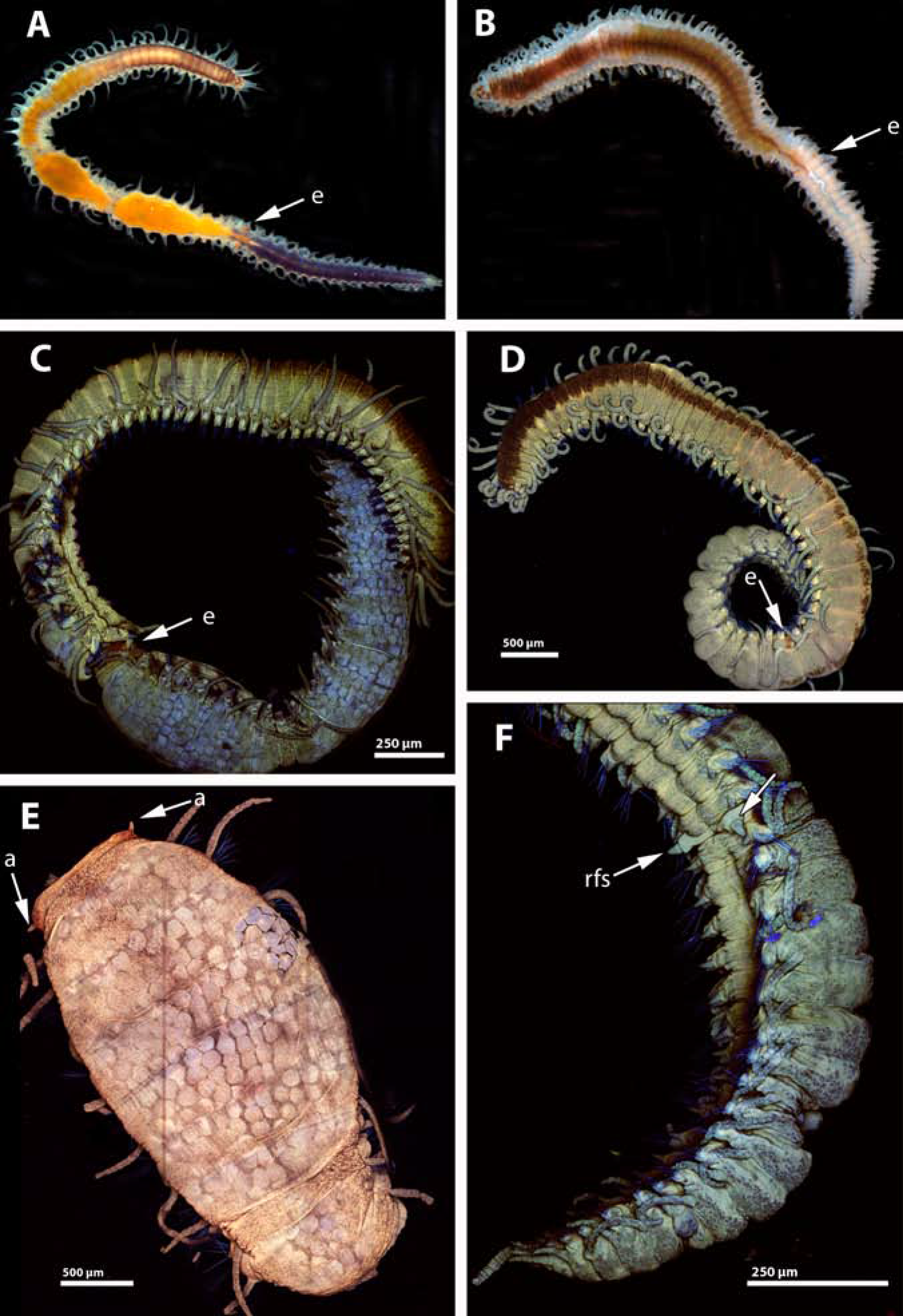
Ligth microscopy pictures of *Syllis magdalena* stolonizing female (A) and male (B). Confocal micrographs of *S. magdalena* stolonizing female (C), male (D), female stolon (E) and male stolons (F). Arrows in A–D pointing to the eyes of stolons (e). Arrows in E pointing to antennae (a). Arrow in F pointing to the regeneration of the final segments in the stock (rfs).

The epithelium of the female and male stolons was columnar, comprised by large epithelial cells (>10 μm in maximum length) with basal non-nucleolated nuclei, and large globular glandular cells with electrondense material (Fig. 3C). In both stolons, below the epithelia, there was a thick layer of muscle fibers, then the germinative epithelium, and finally the digestive epithelium (Fig. 3C–F). The muscle fibers of both female and male stolons presented the regular morphology of muscle fibers of the adults, with a double striation and 25–35 myofilaments and clusters of mitochondria near the tips (Fig. 3C, E). We did not observe the “stolonal” muscle fibers described in *Syllis amica* with the mitochondria towards the middle of the fiber (Wissoq 1967) while attached to the stock. It is possible that the reorganization of the muscle fibers takes place later in the stolonization process, but it is improbable, given that it occurs during head formation in the stolon of *S. amica* (see Wissoq 1967), a process that we observed in *S. magdalena*.

**Figure 3.**
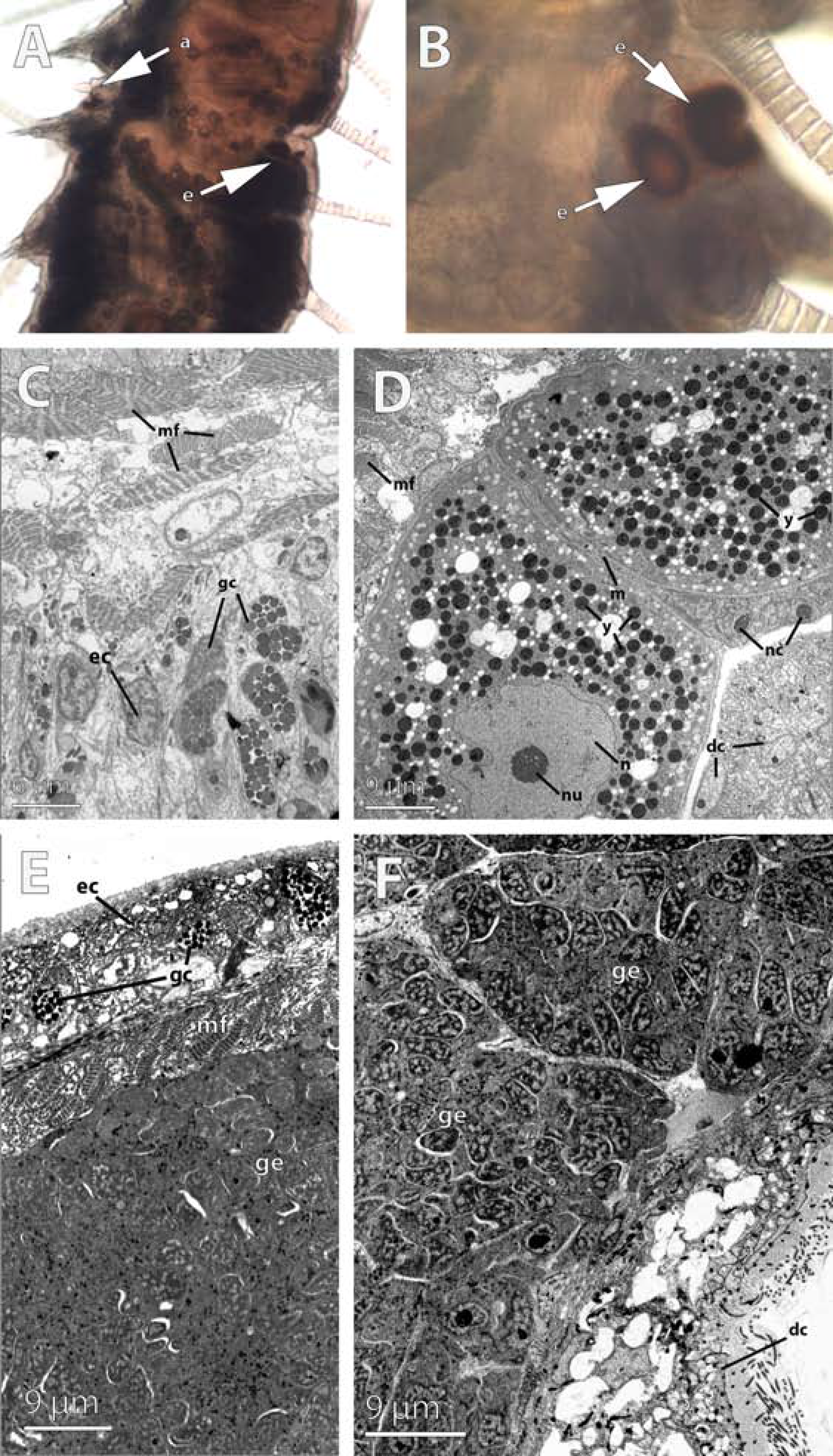
Ligth and electron microscopy pictures of the anterior part of the female and male stolons of *Syllis magdalena*. (A-B) Location of antennae (a) and the two pairs of eyes (e) in the female stolon. (C) Trasmission electron micrographs of the epithelium of the female stolon showing the muscle fibers (mf), granular cells (gc) and epithelial cells (ec). (D) Developing oocytes showing nucleolate (nu) nucleus (n), ooplasm filled with yolk platelets, and microvilli (m) contacting close oocytes. Note the muscle fibers (mf) nurse cells (nc) and the digestive epithelium (dc) surrounding the germinal epithelium. (E-F) Germinal epithelium (ge) in the male stolon. The stolonal epithelium is comprised by a layer of epithelial cells (ec) with interspersed granular cells (gc), and a layer of muscle fibers (mf); spermatogonia develop in the germinal epithelium (gc) below. The digestive cells (dc) lay below the germinal epithelium.

In the female germinative epithelium, large yolky oocytes (50 μm approximately) were surrounded by non-nucleolated nurse cells (Fig. 3D). Oocytes were connected by microvillar processes (Fig. 3D). The male germinative epithelium only contained two large sacs of spermatogonia in the specimens collected (Fig. 3E–F). Spermatogonia (ca. 1 μm in diameter) were densely packed and possess a non-nucleolated nucleus with chromatin condensation processes (Fig. 3E–F). The digestive epithelium was comprised of large (>10 μm in maximum length) convoluted multiflagellated cells (Fig. 3F). We did not observe digestive material in the lumen of the stolon gut (Fig. 3F). There were no differences in the developmental stage of gametes between the anterior and posterior parts of stolons (see also differential expression results).

### General characterization of the de novo transcriptomes

A total 30 libraries derived from somatic and reproductive tissues of non-reproductive (NON-REPRO) and reproductive (REPRO) specimens, were used to assemble de *novo* two transcriptomes, REFSOM and REFTOTREPRO (further details in Material and Methods section). Three somatic parts were chosen for RNA extraction were chosen from all specimens: anterior part (prostomium+first 2 segments), proventricle (all segments containing the proventricle) and final part (pygidium+2 final segments). Two additional fragments from specimens engaged in stolonization (REPRO) were sequenced: the anterior and posterior half parts of the stolons. Assembly statistics of the reference transcriptomes are summarized in Supplementary File S1, alongside read mapping results for each tissue and specimen. The coverage of our assemblies is similar or slightly higher than those in other studies on marine invertebrates (e.g., Meyer et al. 2009; Riesgo et al. 2012; Pérez-Portela et al. 2016).

An overview of the assigned GO terms for each transcriptome (including 3 different levels: cellular component, CC, biological process, BP, and molecular function, MF) and GO enrichment analyses using Fisher’s tests are shown in Supplementary File S2A. The GO enrichment results for the comparisons of both transcriptomes showed 36 GO terms overrepresented in REFSOM related to cellular organization and regulation, metabolism and binding, among others (Supplementary Figure S2B). In contrast, only 8 categories appeared enriched in REFTOTREPRO, mainly related to signaling activity (Supplementary Figure S2C). Interestingly, one of these enriched categories is the activity of G-protein coupled receptors, which bind light-sensitive compounds, pheromones, hormones, neurotransmitters and other ligands involved in secretory processes or cell development, among other functions (e.g., Li et al. 1999; Iversen et al. 2002; Hauser et al. 2006; Asahara et al. 2013). The results of several of these G-protein coupled receptor expression levels on the different tissues and conditions analyzed are discussed below.

### Differential gene expression analyses

#### Pairwise comparisons of somatic tissues (anterior part, proventricle, final segments) between REPRO and NON-REPRO

We detected 792 differentially expressed genes in the comparison between REPRO and NON-REPRO somatic tissues, 494 of them being upregulated in REPRO (178 in females and 316 in males) and 298 in NON-REPRO (Fig. 3 and Supplementary Files S3 and S4). Of these 792 genes, only 292 (~37%) had a BLAST hit and, therefore only the putative annotations for those genes (Supplementary File S4) are discussed below. Among the pairwise comparisons of REPRO and NON-REPRO tissues, the final segments tissue are the ones that showed more differentially expressed genes, was the final segments (Fig. 4C), with 223 differentially expressed in the comparison of female final segments and NON-REPRO final segments (152 upregulated in female) and 460 differentially expressed genes in the comparison of male final segments and NON-REPRO final segments (304 of those upregulated in male). The pairwise comparisons of anterior part and proventricle between reproductive and non-reproductive individuals showed low numbers of differentially expressed genes (Fig. 4A–B). Among those, the largest differences were in the proventricle with 7 differentially expressed genes upregulated in both females and males when compared to non-reproductive, and 20 and 36 differentially expressed genes upregulated in the proventricle of non-reproductive individuals (Fig. 4B).

**Figure 4.**
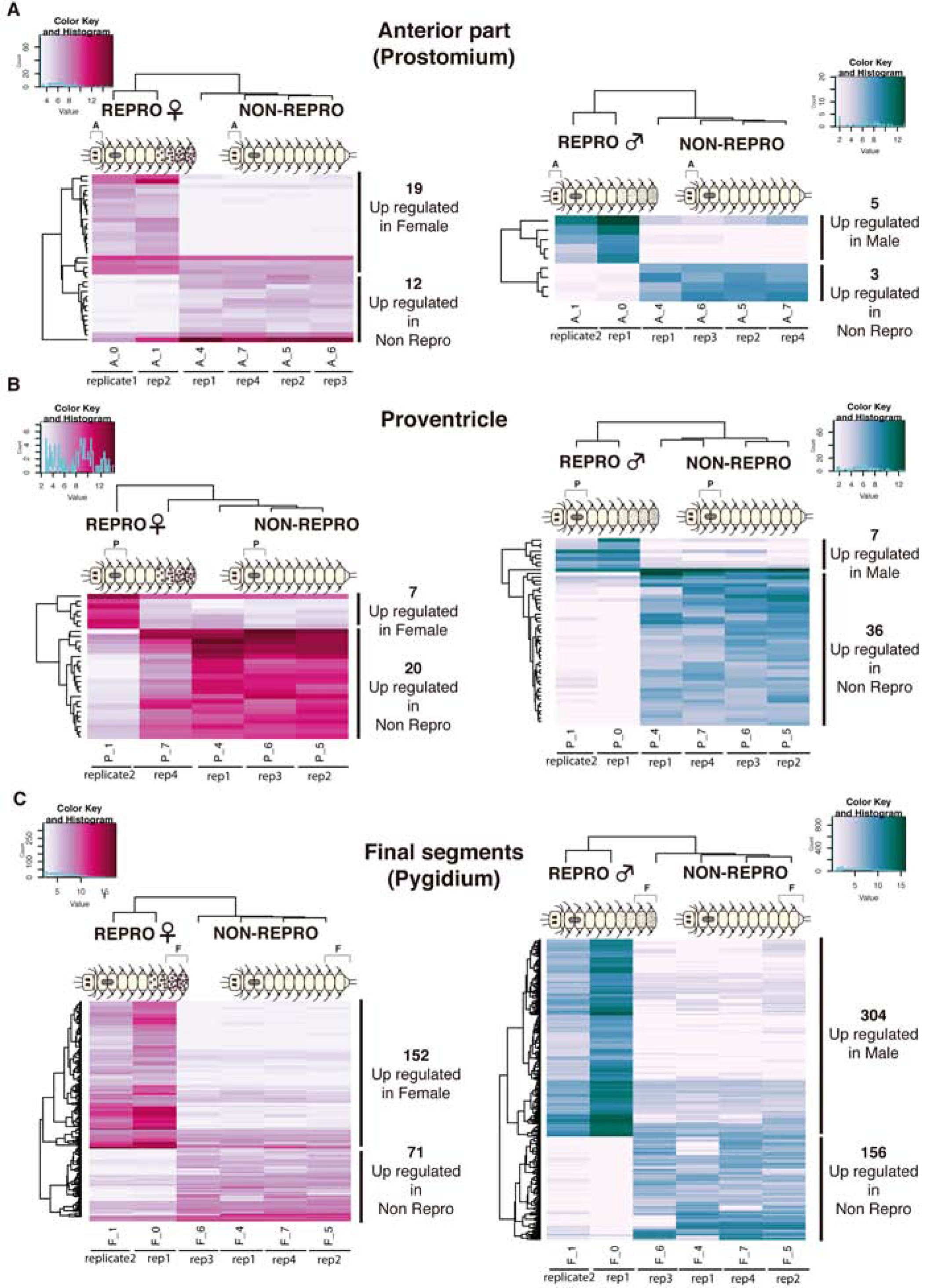
Heatmaps of differentially expressed genes (annotated and not annotated genes) from pairwise comparisons of somatic tissues between reproductive (both female and male) and non-reproductive individuals. Anterior part tissue comparisons (A), proventricle comparisons (B) and final segments comparisons (C). Different colors indicate relative expression levels (see color key and histogram on each). Similarity in expression patterns between genes and individuals is represented by clustering. Abbreviations: anterior part (A), proventricle (P), final segments (F).

In the anterior part and the proventricle of females, the genes upregulated (Supplementary File S4) were related mostly to immune processes (*complement receptor 2*) or food processing (*trefoil factor 2*, *cubilin*, *serine protease 27* and *chitinase*). Similarly, in the male anterior part and proventricle (Supplementary File S4), most genes were involved in nutrient transport (*sugar transporter STL1* and *glycogen phororylase*), as well as development of the nervous system (*tyrosine-protein kinase Src42A*).

Several genes related to gametogenesis were found differentially expressed in the final segments of female and male REPRO individuals compared to NONREPRO (Supplementary File S4), including *vitellogenin* (*Vtg*) and *ovochymase* (*OVCH*) in females, and *testis-specific serine/threonine-kinase* (*TSSK*) in males, which indicates an important role of the final segments during the gametogenesis process in both stolonizing females and males. *Vitellogenin* has been already reported to be involved in annelid gametogenesis, specifically as a yolk precursor (e.g., Hafer et al. 1992), but *OVCH*, an ovary-specific gene involved in egg development of several animals (e.g., Lindsay and Hedrick 1995; Gao and Zhang 2009; Mino and Sawada 2016), is here reported for the first time in annelids. The same occurs for *TSSK*, whose expression, confined almost exclusively to testes, has largely been studied in several mammals (Hao et al. 2004), but never in annelids. Remarkably, two hormone receptors for *relaxin* and *follistatin* were found differentially expressed in the final segments of reproductive females (Supplementary File S4). The insulin-related peptide *relaxin* is important for the growth and remodeling of reproductive tissues during mammal pregnancy (e.g., Gunnersen et al.1995; Hsu et al. 2002) and is active in the ovary and during embryogenesis of zebrafish (e.g., Donizetti et al. 2008, 2010; Wilson et al. 2009). *Relaxin* activity has also been reported in invertebrates, including in the tunicate *Ciona intestinalis* (e.g., Ivell et al. 2005; Olinski et al. 2006), and in the starfish *Asterina pectinifera* (Mita 2013; Mita et al. 2014), where it takes part in oocyte release from the ovary, but this is the first time that it is described in annelids. Likewise, *follistatin*, reported as a follicle-stimulating hormone, with several additional regulatory functions both in reproductive and non-reproductive tissues (Phillips and Kretser 1998), has been already found in the transcriptome of other annelids such as *C. teleta* and *S. lamarckii* (Kenny et al. 2014), but without a particular association with any biological process. In our case, it seems that both *relaxin* and *follistatin* are important during oocyte development in *S. magdalena,* as they are expressed in tissues where oogenesis is taking place before oocytes are transferred into the stolon (see also results section below).

#### Pairwise comparisons of somatic (anterior part, proventricle, final segments) between REPRO females and males

We detected 234 genes with significant expression in the comparison between female and male somatic tissues, 85 of them being upregulated in female (0 in anterior part, 27 in proventricle, 58 in final segments) and 149 in males (only in final segments) (see details in Fig. 5A, B and Supplementary File S5). Of these 234 genes, only 84 (~35%) of transcripts were annotated (Supplementary File S5). No differential expression was found in the comparisons of the female and male anterior parts, and in the proventricle comparisons, we only found differentially expressed genes in the females (Fig. 5A and Supplementary File S5; see discussion below). Similar to the previous comparisons (see above), the somatic tissue sample that showed more differentially expressed genes was the final segments, with 149 genes upregulated in males and 58 in females (Fig. 5B and Supplementary File S5).

**Figure 5.**
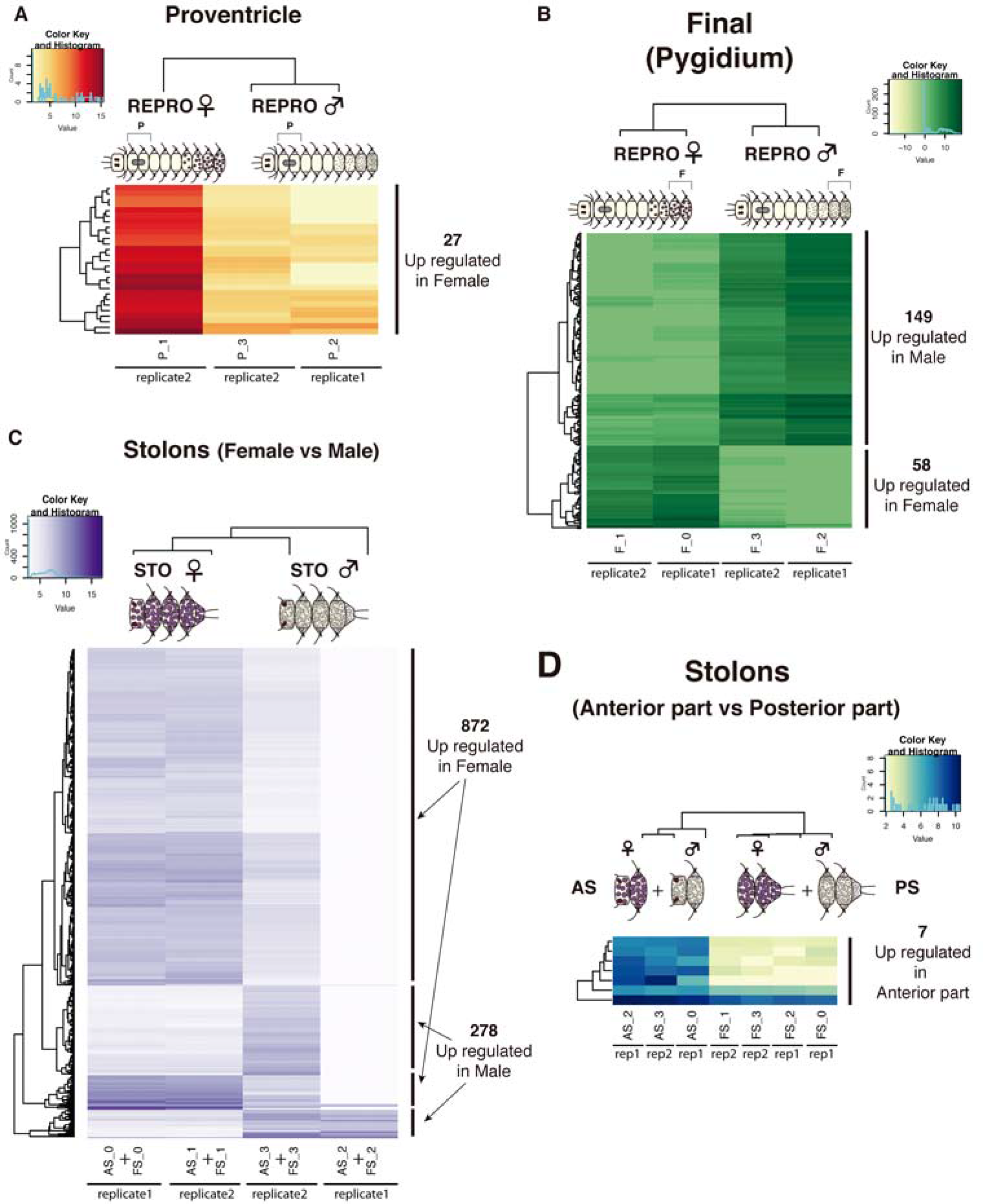
Heatmaps based on differentially expressed genes (annotated and not annotated genes) from pairwise comparisons of somatic tissues between females and males (A, B) and reproductive tissues (stolons) (C, D). Proventricle comparisons (A), final segments comparisons (B), female and male stolons comparisons (C) and anterior and posterior parts of stolons (female and male together) (D). Different colors indicate relative expression levels (see color key and histogram on each). Similarity in expression patterns between genes and individuals is represented by clustering. Abbreviations: anterior part (A), proventricle (P), final segments (F), anterior half of stolon (AS), posterior part of stolon (FS).

As in the previous comparisons (see section above), several gametogenesis-related genes, such as *vitellogenin*, *ovochymase* (*OVOCH*) in females, and *TSSK* in males, were differentially expressed in F (Fig. 5B and Supplementary File S5). In addition, we also found *NOTCH* differentially expressed in F of REPRO males (Fig. 5B and Supplementary File S5). This gene has been reported to have a role in segment formation and adult regeneration in annelids (e.g., Tham and Seaver 2008), and therefore may also be involved in segment formation of stolons and pygidium regeneration of *S. magdalena* (Fig. 2F). However, the *NOTCH* pathway has been also reported to be essential for the correct development of gametes in *D. melanogaster* and mammals (Xu et al. 1992; Hayashi et al. 2001; Murta et al. 2014), and therefore it could also be playing such role during spermatogenesis in *Syllis magdalena*.

Two different transcripts of *ovochymase* were differentially expressed in final segments (*OVOCH1*) and proventricle (*OVOCH2*) female tissues (Fig. 5A and Supplementary File S5). *Ovochymases* are involved in the oogenesis in other invertebrates, where they help avoid self-fertilization and are localized in the vitelline coat of oocytes (Mino and Sawada 2016). In the ascidian *Halocynthia roretzi*, *ovochymase* has a signal peptide, three trypsin-like serine protease domains and six CUB domains (Mino and Sawada 2016). We found 3 *ovochymases* (two DE, *OVOCH1* and *OVOCH2*, and one non-DE, *OVOCH3*) in *S. magdalena*, none of them containing a signal peptide and all containing much less trypsin-like serine protease and CUB domains (Supplementary File S6). The trypsin-like serine protease domain is not exclusive to *ovochymases*, since it also occurs in chymotrypsins (Supplementary File S6), which are digestive enzymes. Given the digestive function of the proventricle in syllids, *OVOCH1* and *OVOCH2* may be performing different functions in *S. magdalena* F and P tissues, respectively. Our molecular phylogeny of ovochymases and chymotrypsins in animals confirmed that *OVOCH1* and *OVOCH3* are homologues of other animal *ovochymases*, whereas *OVOCH2* (the one differentially expressed in the proventricle) is, in fact, a homologue of a mollusk chymotrypsin (Supplementary File S6). *OVOCH1* in *S. magdalena* could be assisting in the maturation of the oocyte, creating an envelope that could further prevent self-fertilization during gamete release in the water column.

#### Pairwise comparisons of stolons between REPRO females and males

We detected 1150 differentially expressed genes in the comparison between reproductive tissues of female and male individuals, 872 upregulated in female stolons and 278 in male stolons (Fig. 5C and Supplementary File S5). This comparison showed the largest differences, with ~75% of genes upregulated in females (872) and ~25% in males (278) (Fig. 5C and Supplementary File S5). In addition, we also compared the anterior and posterior halves of stolons, finding only 7 genes upregulated in the anterior half part (Fig. 5C and Supplementary File S5), most of them related to eye (*rhabdomeric opsin*, *retinal-binding protein*) or brain (*TRPC channel protein*) functioning.

Among the most upregulated Biological Process categories in female stolons, we found Nicotinamide metabolism (Fig. 6). Cells need to accommodate the bioenergetic demands During oogenesis, nicotinate and nicotinamide are essential for organisms as the precursors for generation of the coenzymes NAD+ and NADP+, which are fundamental in redox reactions and carry electrons from one reaction to another, being the pillars of many metabolic pathways. The gene *nicotinamide mononucleotide adenylyltransferase 1-like*, which catalyzes the formation of NAD+, was upregulated in the female stolon when compared to the male stolon (Supplementary File S5). Other metabolic pathways upregulated in the female stolons include both fructose and carbohydrate metabolism, illustrating the high energetic requirements of oogenesis (Fig. 6). In male stolons, the major upregulated process related to the high energetic demands of spermatogenesis is Purine metabolism, a pathway required for nucleotide biosynthesis (Fig. 6). Interestingly, the MAPK cascade (included in the category ‘Styrene catabolism’), which is central to cell proliferation, is upregulated in female stolons (Fig. 6). Similarly, the gene *alpha-1D adrenergic receptor-like*, which also regulates cell proliferation is upregulated in female stolons.

**Figure 6.**
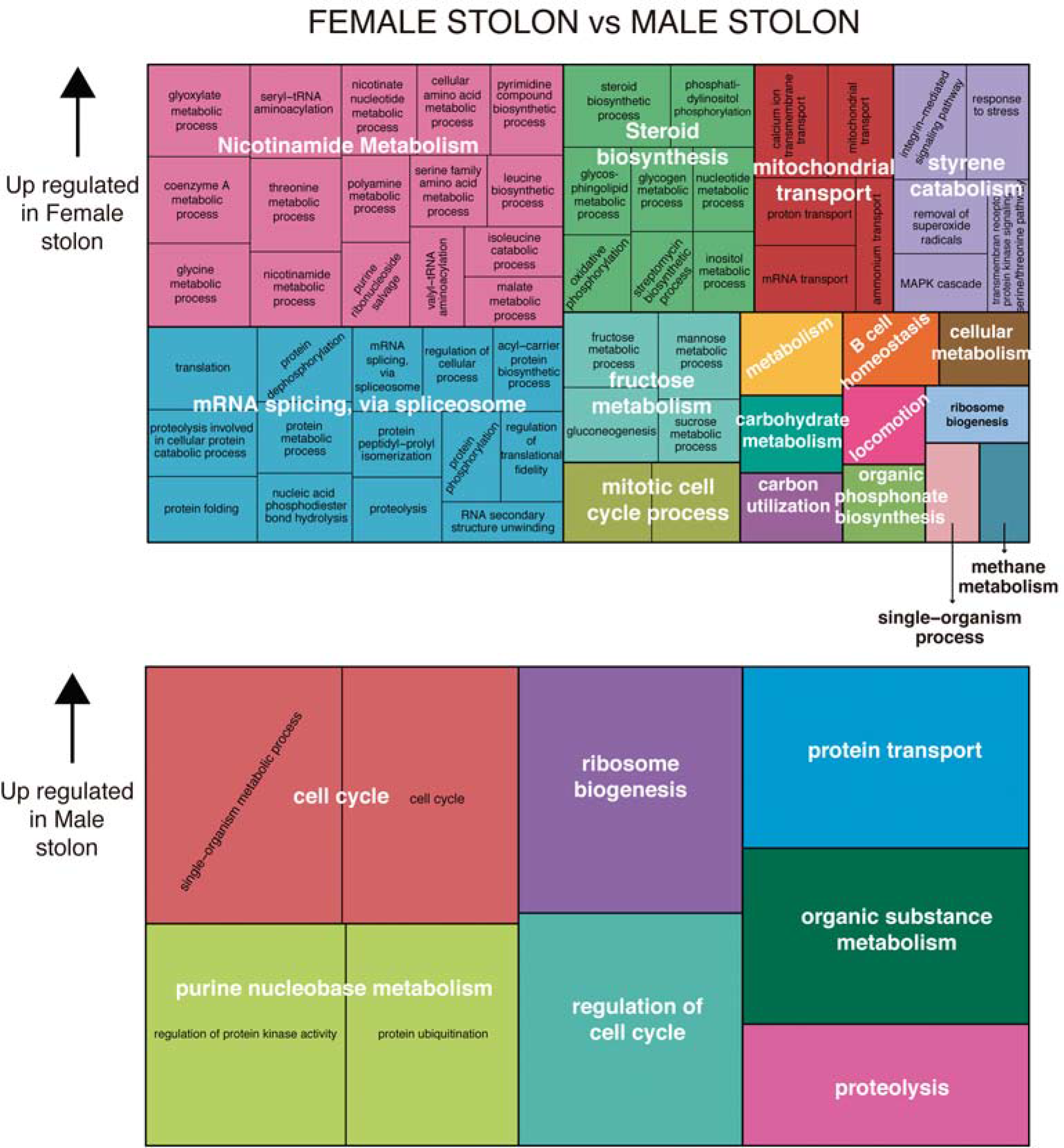
Gene Ontology treemaps for annotated differentially expressed genes in female stolons versus male stolons. The GO terms downregulated in female stolons, are upregulated accordingly in male stolons.

As in the case of final segments (see section above), *Vtg* and *OVOCH* in females, and *TSSK* and *NOTCH* in males, were also differentially expressed in stolons of females and males (Fig. 5C and Supplementary File S5). These results indicate an important role of the stolons in the maturation of gametes, in contrast to what has been traditionally suggested, where the stolons are thought to be only a place to keep and later spread the gametes. However, no genes related to gamete maturation were found to show differentially expressed in the comparison between the anterior and posterior halves of stolon, which suggest that there is no sequential anteroposterior maturation of gametes within the stolons (Fig. 5D and Supplementary File S5), in agreement with our results from the morphological and ultrastructural study.

*Relaxin* was also found differentially expressed in female stolons, reinforcing the hypothesis of its implication in annelid oogenesis and its potential role in the release of oocytes into the water column, as it has been suggested for *relaxin* in *A. pectinifera* (Mita et al. 2014). Other genes involved in gametogenesis of annelids (e.g., Rebscher et al. 2007; Dill and Seaver 2008; Novo et al. 2013) were also found differentially expressed in female stolons (Supplementary File S5), including the member of the DEAD-box helicase protein family, *vasa*. We found two paralogues of the gene *vasa* (the DE *vasa1* and the non-DE *vasa2*) among our transcripts, in contrast to what is found in other annelids that only present one (see Supplementary File S7). While *vasa 2* grouped with all *vasa* orthologs obtained in annelids, *vasa 1* branched out from the annelids and appeared basal to other vasa orthologs from metazoans (Supplementary File S7), being more similar to *ATP-dependent RNA helicase vasa-like* proteins in arthropods than to *vasa* proteins of annelids when blasted. These results may suggest that different paralogues may be performing different functions in *S. magdalena* (Supplementary File S7). While *vasa2* could be playing a role in the female germline determination localized in the oocytes of *S. magdalena*, *vasa1* could be participating in the maintenance of totipotency of the stem cells (Juliano and Wessel 2010), although *ATP-dependent RNA helicase vasa-like* proteins are also known to be involved in oogenesis. Interestingly, we also found the category Steroid biosynthesis upregulated in female stolons (Fig. 6). In addition, our study shows the upregulation of the gene *hydroxysteroid dehydrogenase 2 isoform X2*, that could potentially mediate steroid hormone metabolism (Seckel and Walker 2001), and suggests hormonal control over the final stages of stolonization in *S. magdalena*.

In male stolons, most of the upregulated genes were involved in the construction of the flagellar apparatus (Inaba 2011), including *dyneins*, *cilia-and the flagella-associated proteins*, *ropporin*, *radial spoke 3*, and *kinesins*). This is unsurprising, given the presence of sperm in these tissues, but is an excellent positive control.

### Hormonal control of stolonization

Since methyl farnesoate (MF) was discovered to be produced by mandibular organs of numerous crustaceans, this form of the insect Juvenile Hormone (JH III), has been commonly considered as the crustacean equivalent of insect JH (Laufer and Biggers 2001; Miyakawa et al. 2014). Comparably to JH in insects, MF regulates many aspects of crustacean physiology, including reproduction (Xie et al. 2016). In this context, MF is more actively synthetized by females during vitellogenesis, and higher levels of MF are associated with large reproductive systems and aggressive mating behavior in males of the spider crab *Libinia emarginata* (Laufer et al. 1992). In the annelid *C. teleta*, exogenous extracts of MF were found to affect larval metamorphosis and settlement (Laufer and Biggers 2001), and MF has been recently demonstrated to be directly involved in *P. dumerilii* regeneration and female sexual maturation (Schenk et al. 2016). This latter study not only showed that the decrease of MF levels in the brain induces reproduction and suppresses regenerative capacities in *P. dumerilii*, but it also reported an ortholog of the MF receptor of arthropods (*bHLH-PAS-domain-containing transcription factor methoprene-tolerant receptor*, *MTr*) in the eleocytes (coelomic cells that synthesize yolk via production of *Vtg* protein), demonstrating that this hormone is not restricted to arthropods, as it was assumed (Schenk et al. 2016). Since detection of MF is not possible in RNAseq data, in order to assess whether *S. magadalena* could use a similar molecular signal to determine when to divert resources from somatic functions to reproduction, we investigated if *S. magdalena* also possessed an ortholog of *MTr*, identified as the arthropod and lophotrochozoan sesquiterpenoid receptor (e.g., Konopova and Jindra 2007; Miyakawa et al. 2013; Jindra et al. 2015; Schenk et al. 2016). In our *de novo* transcriptomes, we identified two transcripts encoding bHLH-PAS-domain-containing transcription factor that showed strong similarity to *P. dumerilii MTr*. In fact, our molecular phylogeny of *MTr* revealed that the *S. magdalena* ortholog is closely related to *MTr* orthologs of *P. dumerilii* and *C. teleta* (Fig. 7A). In agreement with Schenk et al. (2016), our results also confirmed that annelid *MTr* is clearly an ortholog of insects and crustaceans *MTrs* (Fig. 7A). These findings allow us to suggest that MF may be one of the hormones responsible for syllid stolonization. If the MF is involved in syllid reproduction, we would expect to find differences in the levels of expression of MF receptors (*MTr*) among the stolonizing and non-stolonizing syllid samples (higher in the latter), similar to what has been reported during oocyte maturation and male reproductive behavior in crustaceans and other annelids (e.g., Laufer et al. 1992; Schenk et al. 2016). Surprisingly, higher expression levels (albeit not statistically significant) of *MTr* were found only in anterior and posterior tissues of female, therefore REPRO, individuals (Fig. 7B), but not in the NON-REPRO specimens, as it might be expected (Schenk et al. 2016). Thus, in contrast to what was found in *P.dumerilii* but similar to what has been reported for arthropods, an increase in MF (or a similar putative sesquiterpenoid) is necessary to initiate the reproductive process in stolonizing syllids (Fig. 7B) (Laufer et al. 1992; Gäde et al. 1997; Wyatt 1997; Hansen et al. 2014). The fact that the differences between conditions are not statistically significant can be explained because the NON-REPRO specimens were collected only one week before the beginning of the stolonization process, and therefore they might have already entered the initial stages of reproduction without visible morphological changes. On the other hand, as in the case of *A. marina* (e.g., Pacey and Bentley 1992), it is also possible that a non-identified hormone, sesquiterpenoid or otherwise, is orchestrating the important metamorphic changes that occur during syllid stolon development, similarly to what MF and JHs do in arthropods (e.g., Hui et al. 2010; Maruzzo et al. 2012; Wen et al. 2015). However, the presence of sesquiterpenoids is further suggested by other DE gene results, as discussed further below.

**Figure 7.**
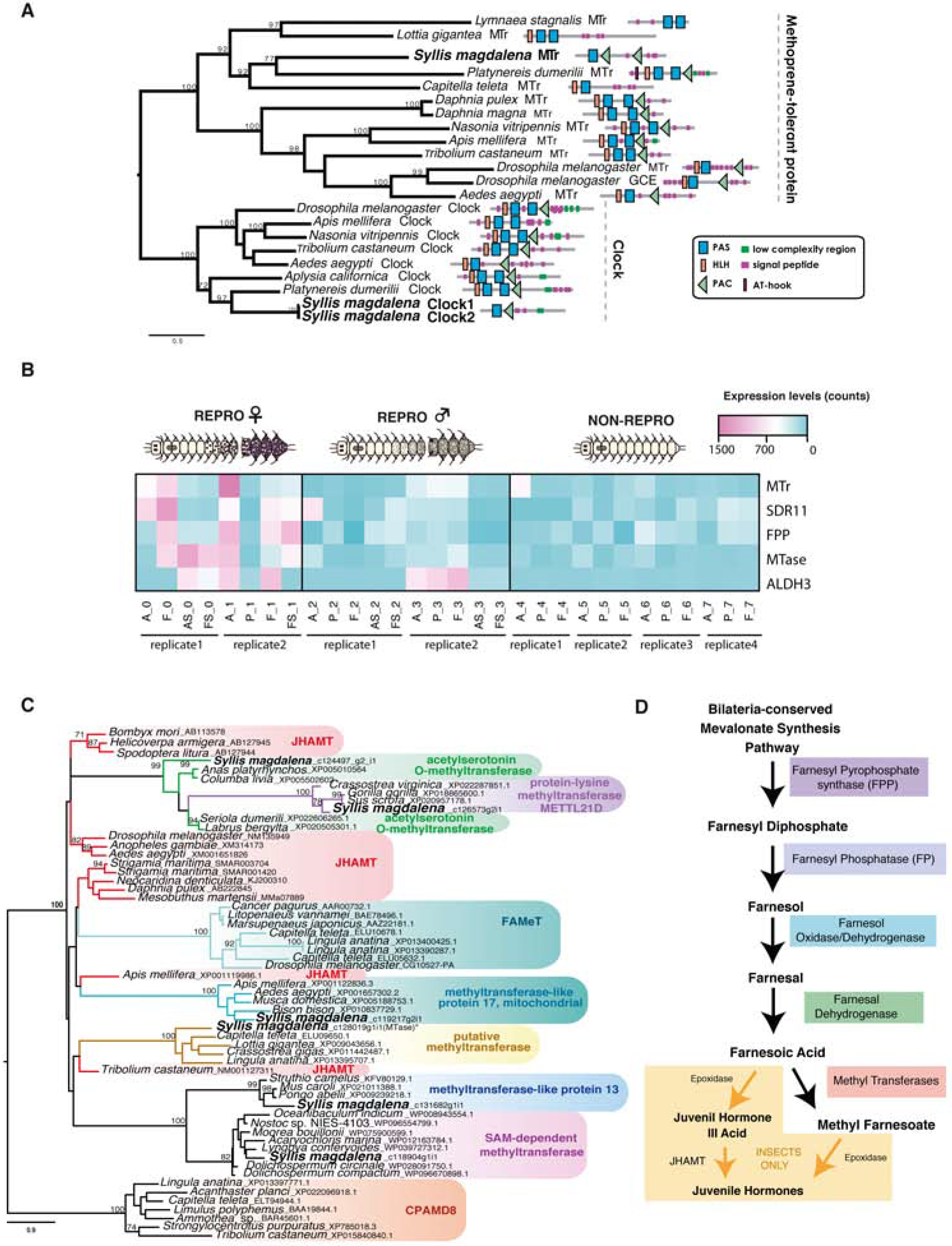
(A) Phylogenetic reconstruction of the protein alignment for methoprene-tolerant receptor (MTr) found in our samples. (B) Heatmap showing the relative levels of expression in the different tissues and conditions analyzed of the transcripts that may be involved in the synthesis of the neurohormone *methyl farnesoate* (MF): MTr, *Farnesol oxidase/dehydrogenase* (SDR11), *Farnesal dehydrogenases* (ALDHE3), the differentially expressed transcript *Farnesyl pyrophosphate synthase* (FPP) and putative *methyl transferase* (Mtase). (C) Phylogenetic reconstruction of the differentially expressed *MTases* in the female stolon. (D) Synthesis pathway of MF and JH in arthropods. Abbreviations: anterior part (A), proventricle (P), final segments (F), anterior half of stolon (AS), posterior part of stolon (FS).

Interestingly, other neurotransmitter receptors were found to be upregulated in the posterior end of NON-REPRO specimens: *dopamine receptor* (*DAr*), belonging to the large family of G-protein coupled receptors, was downregulated in the final segments of females, and *serotonin transporter* (*SERT* or *5-HTT*), which terminates the action of serotonin, was downregulated in the final segments of males (Supplementary File S5 and Fig. 8A). Our molecular phylogeny corroborates that these proteins are orthologs of the *C. teleta DAr type 2* (*DAr2*, Fig. 8B) and *C. teleta* and *Helobdella robusta SERT* genes (Fig. 8B). Dopamine (DA) and Serotonin (SER) are biogenic amines that act as a neurotransmitters and hormones, regulating an array of important physiological functions both in vertebrates and invertebrates (e.g., Winberg et al. 1997; Neckameyer, 1998a; Gingrich et al. 2000; Wicker-Thomas and Hamann 2008; Dufour et al. 2010; Giang et al. 2011). In *D. melanogaster* DA and SER control a wide range of behavioral processes such as circadian rhythms, sleep, mating behavior, learning or aggression (e.g., Nichols 2007; Giang et al. 2011) and also stimulate fertility and female receptivity (Neckameyer 1998b; Marican et al. 2004). In *C. elegans*, male mating behavior and egg deposition are also induced by DA and SER (Sulston et al. 1975; Weinshenker et al. 1995; Dempsey et al. 2005). In addition, both hormones have been reported to be involved in larval metamorphosis in cnidarians, molluscs, and echinoderms (Couper and Leise 1996; McCauley 1997; Matsuura et al. 2009). In annelids, dopaminergic and serotonergic systems have been found in several species (Grothe et al. 1987; Dietzel and Gottmann 1988; Schlawny et al. 1991; Spörhase-Eichmann et al. 1998; Krajniak and Klohr 1999; Zaccardi et al. 2004; Lawrence and Soame 2009; Helm et al. 2014; Rimskaya-Korsakova et al. 2016; Verasztó et al. 2017; Bauknecht and Jékely 2017). However, the participation of DA and SER in annelid reproduction has only been demonstrated in a handful of studies. Although it was thought that DA played an important role in sexual differentiation in *Ophryotrocha puerilis* (Grothe et al. 1987; Pfannenstiel and Spiehl 1987; Grothe and Pfannenstiel 1986), it was later demonstrated that the catecholaminergic system of this species was involved in mechano-and/or chemoreception (Schlawny et al. 1991). In contrast, both SER and DA in nereids seem to have a positive effect on oocyte development, the first by directly inducing their maturation and the second by switching off the action of the juvenile hormone (Lawrence and Soame 2009). Similarly, in the decapod *Penaeus merguiensis* SER induces ovarian maturation through MF production (Makkapan et al. 2011). In this sense, increased levels of both hormones, as indicated by the upregulation of their receptors and/or transporters (*DAr* and *SERt*) just before the beginning of stolonization (NON-REPRO individuals), could be the stimulus required to initiate oocyte and sperm development during syllid stolonization, with a decrease in the levels afterwards during the course of gametogenesis. In addition to this suggested putative direct role in gametogenesis *per se,* DA could also be the putative hormone in the brain and/or proventricle inducing the production of MF (or other sesquiterpenoid) to regulate stolonization in *S. magdalena*, as found for DA and the JH of nereids and decapods (Lawrence and Soame 2009; Makkapan et al. 2011). Our results thus indicate a possible role of several hormonal factors in the sexual differentiation of stolons, in agreement with previous studies (Franke 1980; Heacox and Schroeder 1982).

**Figure 8.**
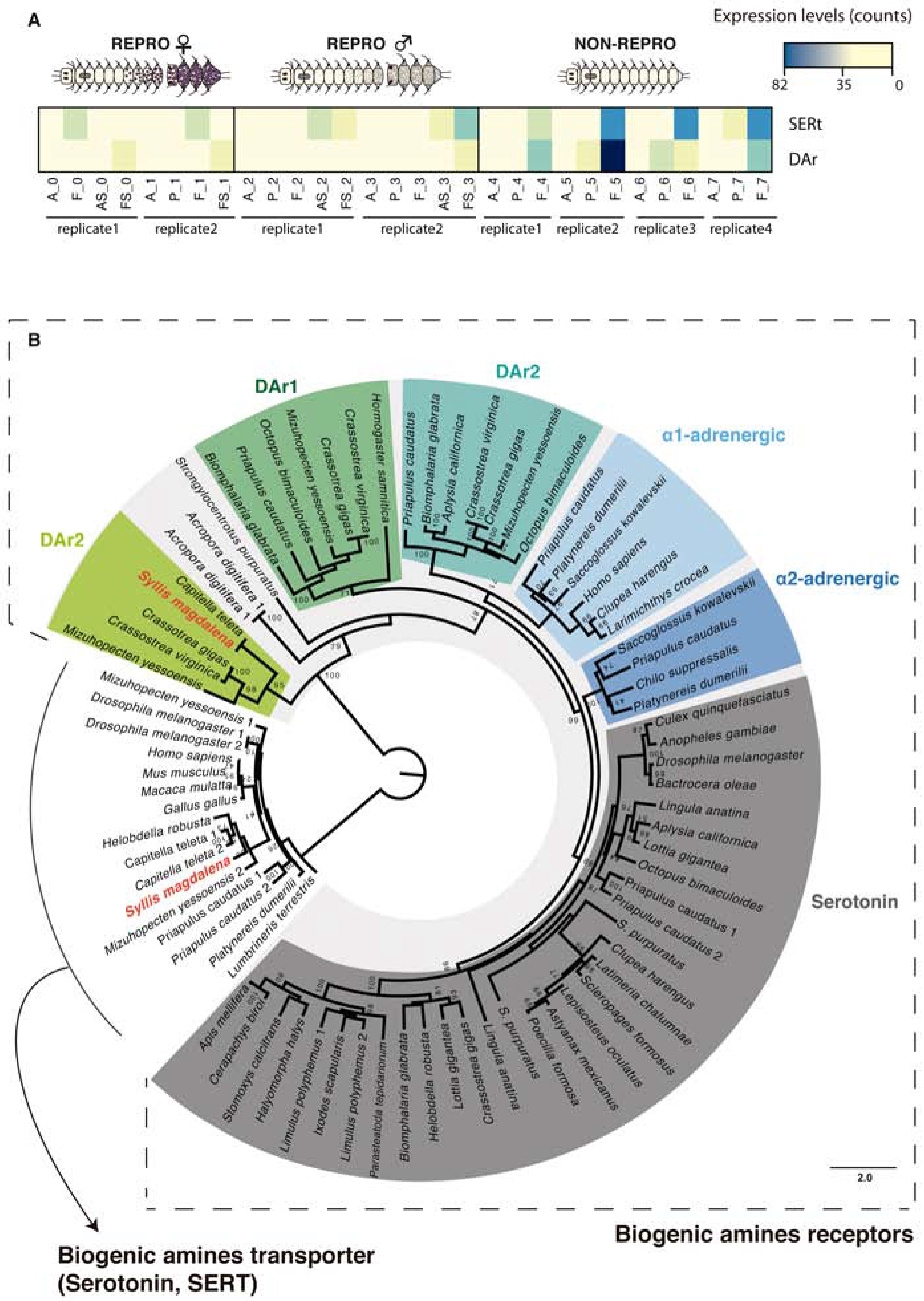
Phylogenetic reconstruction (A) and heatmap of relative levels of expression in all the tissues and conditions (B) of the genes *dopamine receptor* (*DAr*) and *serotonin transporter* (*SERT*). Abbreviations: anterior part (A), proventricle (P), final segments (F), anterior half of stolon (AS), posterior part of stolon (FS).

In addition, if DA and SER were the neurohormones regulating stolonization in syllids, our results do not support the traditional view in which male stolons differentiate autonomously and female stolons differentiate upon hormone release by the male stolon (Franke 1999). We found upregulation of the receptors of these two neurohormones in both female and male individuals at the beginning of stolonization. DA and SER have been reported to be under the influence of photoperiodic and circadian rhythms, which are essential for synchronizing several processes in animals (Andretic and Hirsh 2000; Doyle et al. 2002; Lawrence and Soame 2009). Therefore, we suggest that that both female and male stolon differentiation are triggered by environmental cues regulating the production of DA and SER. As in other annelids, the main external signals that may be controlling the synchronicity of the reproductive period in syllids are light and seawater temperature (e.g., Franke 1986b). In the Adriatic Sea, the breeding season of *Syllis prolifera* is restricted from late March to early October, when the temperature ranges from 14°C–19°C, and there are around 12–13h of light per day (Franke 1986b). Similar results were observed in *S. magdalena*, which seems to breed during the southern hemisphere summer (see sampling methods) with a mean seawater temperature around 15°C and around 13h hours of light per day.

In addition to steroid hormone control, we found some differentially expressed genes in the female stolons, potentially involved in the production of pheromones (specifically the sesquiterpenoid MF; see section above): *Farnesyl pyrophosphate synthase* (*FPPS*) and several *methyl transferases* (*MTases*) (Fig. 7B, C and Supplementary File S5), which could synthetize sesquiterpenoids similar to MF and JHIII in arthropods (e.g., Tobe and Bendena 1999; Hui et al. 2010). Specifically, *FPPS* is required at the beginning of the process to catalyze the reaction, generating Farnesyl Diphosphate, the raw material for sesquiterpenoid production, which is then transformed into Farnesol, then Farnesal (via the *Farnesol oxidase/dehydrogenase*, *SDR11*), later into FA (through *Farnesal dehydrogenases*, *ALDHE3*), and, in the canonical pathway, finally into MF in crustaceans (through *Farnesoic acid methyl transferase*, FAMeT), or into JH in insects (through an *epoxidase*, *FAMeT* and *Juvenile hormone acid O-methyltransferase*, *JHAMT*) (e.g., Hui et al. 2010) (Fig. 7D).

Following Schenk et al. (2016) and given our results (including those for *methoprene tolerant receptor*, above), a similar pathway seems to occur in annelids, with the synthesis of some form of sesquiterpenoid regulating reproduction, as occurs in arthropods (Xie et al. 2016). In fact, our phylogenetic results confirmed that the differentially expressed transcripts annotated as *FPPS* and of a variety *MTases* (Figs. 7C and Supplementary File S8) are legit orthologs, and thus the beginning and end of the synthesis cascade, and the likely bottleneck, are differentially expressed. In addition, orthologs of *FPP*, *SDR11* and *ALDHE3* of spiralians were cleary found in our samples (Supplementary File S9), although these are not differentially expressed themselves. These differentially expressed *MTases* are of a variety of annotations, with some possessing homologs across the Bilateria. None possess clear homology to known arthropod *FAMeT* or *JHAMT* sequences. However, all could potentially be performing a similar role *in vivo*, and one apparent Spiralia novelty is present, which we posit as an excellent candidate for future functional investigation.

However, despite this persuasive circumstancial evidence, we still cannot confirm that the final product of this biosynthetic pathway in *S. magdalena* is MF or another sesquiterpenoid, until functional analyses are performed to test this hypothesis. Besides the putative involvement of sesquiterpenoids in the beginning of syllid stolonization, which is reinforced by the high expression of *SDR11* and *ALDH3* in somatic tissues of both male and female individuals (Fig. 7B), it seems that in our case it may also affect later stages, since *FPPS* and *MTases* are differentially expressed in female stolons (Supplementary File S5). Thus, the increase of MF levels could also be regulating the *vitellogenin* levels necessary for yolk formation, as it commonly occurs with JH in arthropods (Laufer et al. 1992; Gäde et al. 1997; Wyatt 1997; Hansen et al. 2014). In fact, the overexpression of this hormone in stolons could be the triggering signal for the stolon release from the stock. We did not find any enzyme necessary to synthetize hormones or neuropeptides differentially expressed in the male stolons, which might indicate that the synchronicity in the release of female and male stolons might be directly controlled by the female via the production of MF, as it has been also reported during spawning in *A. marina* (Hardege and Benteley 1997).

In addition, as discussed above, MF production has been shown to be influenced by external stimuli (e.g., Shin et al. 2012; Girish et al. 2015; Toyota et al. 2015), which could trigger the stolonization process simultaneously in syllid species according to the traditional hypothesis (e.g., Franke 1999). One of these external stimulus is ambient light variation, which is detected via photosensitive pigments such as opsin proteins and represents a common mechanism mediating the synchronization of gamete release or spawning in a variety of marine invertebrates (Kaniewska et al. 2015; Siebert and Juliano 2017). We have identified several *opsin* homologs in *S. magdalena*, including a *rhabdomeric opsin* previously characterized in other annelids (e.g., Arendt et al. 2004; Randel et al. 2013; Gühmann et al. 2015), that was found differentially expressed in the anterior part of stolons (Supplementary File S5), but not in the anterior part of the stock. Our molecular phylogeny including all *opsins* found in *S. magdalena* (Fig. 9A) revealed that the differentially expressed *rhabdomeric opsin* (*r-opsin 5*) and two other non-differentially expressed *opsins* (*r-opsin 3* and *4*) are homologues of the *P. dumerilii opsin* found in larval eyes (Arendt et al. 2002). Differences on expression levels among tissues and conditions were observed in the different opsins found in our samples (Fig. 9B), which suggest several roles of opsins at different stages of syllids development, as it has been already stablished in other marine annelids (e.g. Arendt et al. 2004). Specifically, the upregulation of *r-opsin 5* in the anterior part of the stolons, where the stolon eyes are located (Fig. 2A–B, 3A–B) suggests that this *opsin* copy in particular might be responsible for detecting the light changes that would trigger MF production, and the subsequent synchronous stolon release and spawning in *S. magdalena*. A similar mechanism has been recently demonstrated in the hydrozoan jellyfish *Clytia hemisphaerica*, in which spawning is mediated by oocyte maturation-inducing neuropeptide hormones (MIHs), whose release is triggered as a response to blue-cyan light detected by a gonad photosensory *opsin* (Artigas et al 2018).

**Figure 9.**
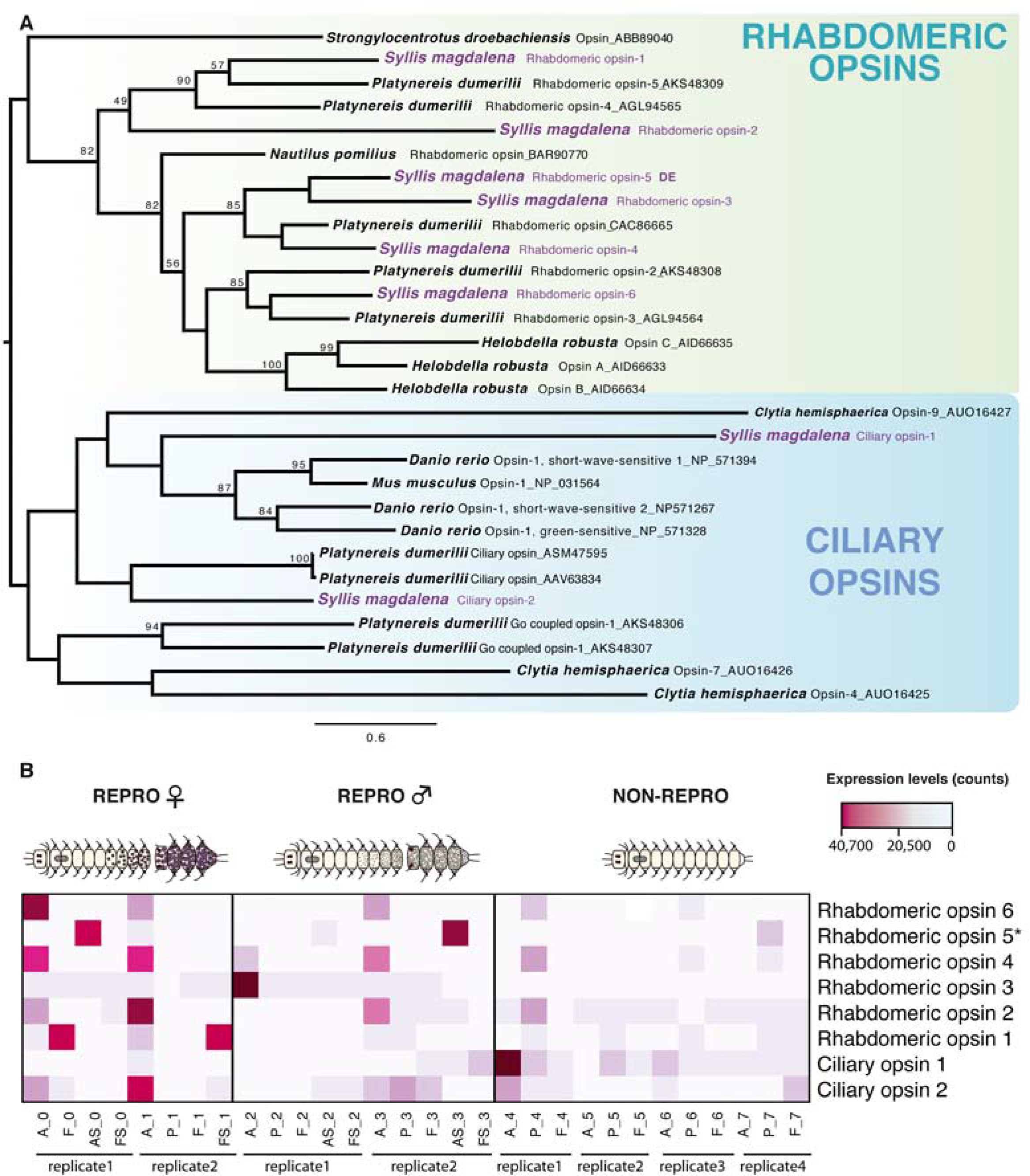
Phylogenetic reconstruction of the protein alignment for the different *opsin* genes (*rhabdomeric* and *ciliary*) found in our samples (A) and levels of expression of all of them in the different tissues and conditions analized (B). *Rhabdomeric opsin 5* appeared differentially expressed in the anterior part of stolons. Abbreviations: anterior part (A), proventricle (P), final segments (F), anterior half of stolon (AS), posterior part of stolon (FS).

### Conclusions

Using Illumina RNA-seq data, we provide the first transcriptomic characterization of the reproductive process in a species of the family Syllidae. Here, we performed a series of pairwise comparisons of the gene expression patterns in different tissues and conditions that allowed us to identify the molecular mechanisms underlying the stolonization process of *Syllis magdalena*. We found an array of differentially expressed genes involved in immune response, neuronal development, gametogenesis, cell proliferation, and steroid metabolism playing different roles in the reproductive process of *S. magdalena*. Among the most striking results of our study was the continuous gamete maturation occurring in both the final segments and the stolons and the hormonal regulation of the reproduction. Thus, following previous hypotheses proposed for other annelids, including syllids (e.g., Franke and Pfannenstiel 1984; Pacey and Bentley 1992; Franke 1999; Lawrence and Soame 2009; Schenk et al. 2016), we suggest a multihormonal model for the control of syllid stolonization, influenced by environmental signals affecting the anterior part (prostomium) and proventricle of the animal, as it was traditionally hypothesized (e.g., Franke 1999), but also influencing the posterior end of the animals (and thus, the gonads) (Fig. 10). When the breeding season approaches, both DA and SER levels increase triggered by photoperiod and circadian rhythms (Andretic and Hirsh 1999; Lawrence and Soame 2009) and they produce a direct influence on the gonads of pre-reproductive individuals (upregulation of DAr/SERt in final segments of NON-REPRO), initiating the gamete production (Fig. 10A, B). The increase of DA and SER could also positively regulate the production of the putative brain and/or proventricle hormones (such as MF or similar), as in several other invertebrates (Couper and Leise 1996; McCauley 1997; Matsuura et al. 2009) regulating the gamete production (and the consequent metamorphosis to produce stolons), as observed in crustaceans and insects (e.g., Shin et al. 2012; Girish et al. 2015; Toyota et al. 2015). At this point, a variety of other hormones and proteins, such as *Vtg*, *OVCH*, *relaxin*, *follistatin* and *TSSK*, play their role in the correct development of gametes (Fig. 10B) until maturation is completed. During gamete and stolon maturation, high levels of MF may be required for yolk formation (upregulation in female stolon of *Vtg, FPPS and MTases*), and the presence of MF could additionally trigger stolon release from the stock as a response to external stimuli (as indicated by the upregulation of photosensitive *r-opsins*) (Fig. 10C). We also suggest that the synchronicity of the stolon and gamete release may not only be mediated by exogenous factors such as light and water temperature, but also by chemical cues provided by the female stolons, as demonstrated in other annelids (Hardege and Bentley 1997).

Overall, our results illuminate the process of stolonization in syllids, improving our understanding of how some putative hormones and gametogenesis-related genes regulate the reproduction in stolonizing syllids. However, the transcriptomic approach adopted here does not allow us to locate the specific expression of these genes, and further functional studies are needed to provide a more complete overview of the expression patterns and the proper functioning of specific pathways during reproduction in *S. magdalena*. In addition, RNAi experiments to inhibit the expression of G-protein coupled receptors and other hormones and neuropeptides would provide promising routes to understand their role during stolonization in syllids, allowing us to elucidate once and for all how these annelids delegate sex to their stolons.

**Figure 10.**
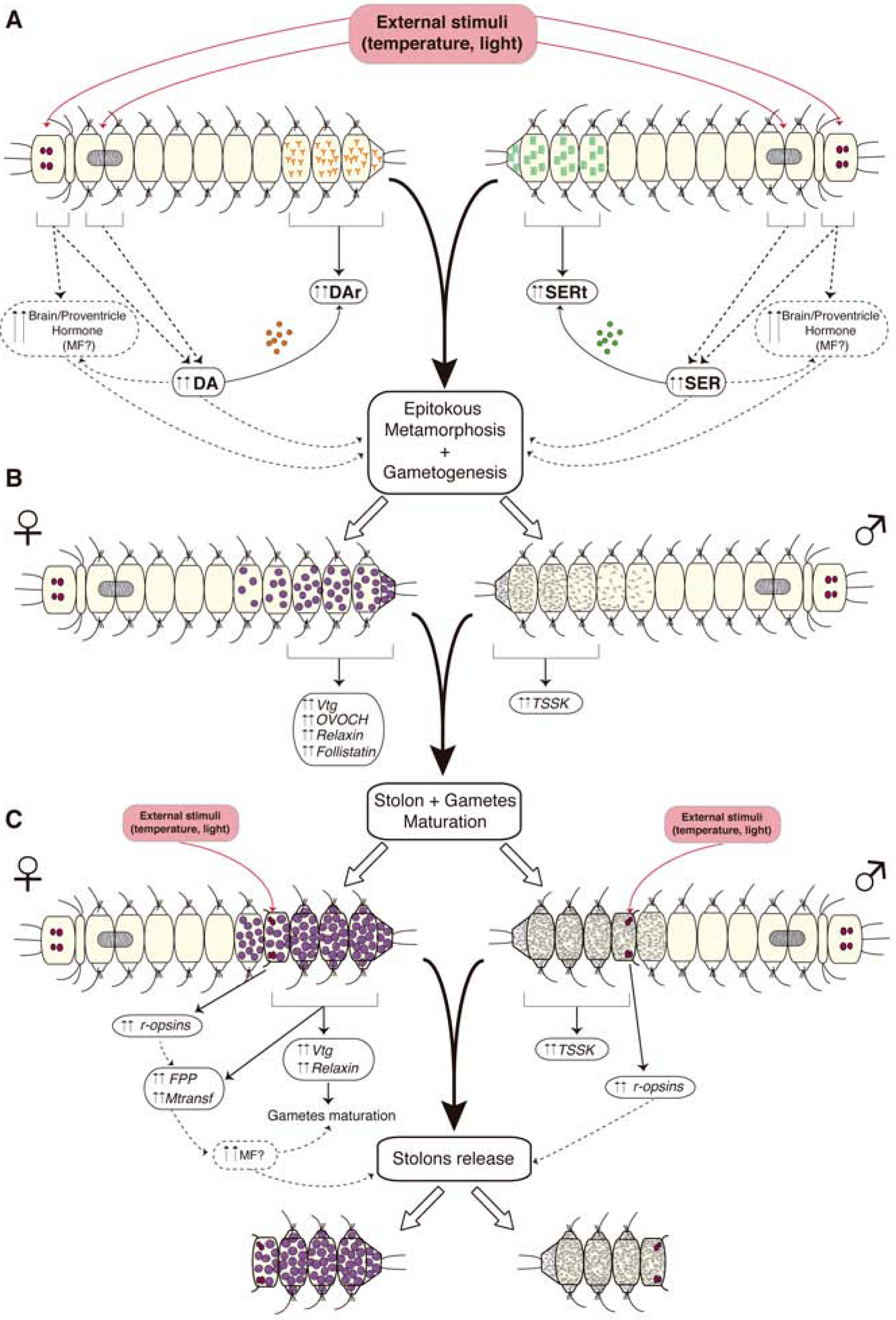
Multihormonal model proposed for the control of syllid stolonization.

## Material and methods

### Sample collection and preservation

Eight individuals of *S. magdalena* were collected in intertidal algal turfs of *Ulva rigida* and *Perumytilus purpuratus* beds, in Las Cruces, Central Chile (S 33° 30’ 06”, W 71° 37’ 55”) in January 2014. Four specimens were collected during full moon, two of which were developing female stolons and the other two male stolons (REPRO specimens); the other four specimens were sampled before the full moon and were not engaged in reproduction (NON-REPRO specimens). All samples were immediately fixed in RNA*later* and stored at −80 °C until RNA extraction. Two additional male and female stolons were preserved complete in 2.5% glutaraldehyde in 0.4M PBS for electron and confocal microscopy.

### Confocal and transmission electron microscopy

Whole specimens preserved in 2.5% glutaraldehyde were mounted in slides to obtain images of auto-fluorescent tissues during stolonization with a Nikon Eclipse upright with A1-Si confocal microscope at the Image Analysis Center (IAC) of the Natural History Museum of London. No stain was applied, but images were obtained in DAPI 488, 555 and 647 channels, under gentle laser excitation. For transmission electron microscopy (TEM), specimens fixed in 2.5% glutaraldehyde were later postfixed in 1% osmium tetroxide and rinsed twice in PBS before dehydration with an increasing series of acetone (from 50% to 100%). Samples were further embedded in epoxy resin, serially sectioned with an ULTRACUT ultramicrotome at 64 nm, post-stained with uranyl acetate and lead citrate, and observed with a JEOL JEM1010 microscope at the Serveis Científico-Tècnics (SCT) at the Universitat de Barcelona and at the Servicio Interdepartamental de Investigación (SIDI) of the Universidad Autónoma de Madrid.

### RNA extraction

Each tissue sample was transferred to a microcentrifuge tube containing 500 μL of TRIzol (Invitrogen), and ground with a RNase-free plastic pestle to break down the tissue, and isolate RNA and DNA. Then, another 500 μL of TRIzol and 10 μL of glycogen were added. After 10 min incubating the mixture at room temperature (RT), 100 μL of the RNA-isolating reagent bromochloropropane (BCP) was mixed in by vortexing. After 10 min incubation at RT, samples were centrifuged at 16,000 relative centrifugal force (rcf) units for 15 min at 4 °C to separate the solution into 3 layers. The upper aqueous layer, which contained total RNA, was recovered and mixed with 500 mL of isopropanol, and incubated at –20 °C overnight. Afterwards, the sample was centrifuged at 16,000 rcf for 15 min at 4 °C, and the supernatant was removed. Total RNA precipitation was performed by washing the remaining pellet twice by adding 1 mL of 75% ethanol and centrifuging it at 16,000 rcf at 4 °C for 5 min. The dried pellet was eluted in 100 μL of RNA Storage solution (Invitrogen). mRNA purification was performed with a Dynabeads mRNA Purification Kit (Invitrogen), following manufacturer’s instructions. After incubation of total RNA at 65 °C for 5 min, the samples were incubated for 30 min with 200 mL of magnetic beads in a rocker and washed twice with washing buffer.

Thirteen μL of 10 mM Tris-HCl were added to the eluate, and the mixture was incubated at 80 °C for 2 min. The supernatant was immediately transferred to a 0.5 mL microcentrifuge tube and stored at −80 °C. Quality of mRNA was measured with a pico RNA assay in the Agilent 2100 BioAnalyzer (Agilent Technologies). Quantity was measured with an RNA assay in a Qubit fluorometer (Life Technologies). Further details about RNA prep protocols can be found in Fernández et al. (2014).

### cDNA Library Construction and Next-Generation Sequencing

cDNA libraries were constructed from extracted mRNA in the Apollo 324 automated system using the PrepX mRNA 8 Protocol Kit (IntegenX) set to 200 base pairs (bp) and stranded mRNA, under the Library Prep Illumina setting. A Polymerase Chain Reaction (PCR) was run to amplify cDNA libraries, using the KAPA Library Amplification Kit. PCR was run as follows: denaturation (45 sec at 98 °C), cycling (15 sec at 98 °C, 30 sec 60 °C, and 15 sec at 72 °C, for 16 cycles), and final extension (1 min at 72 °C). During the PCR process, the samples were marked with a different index to allow pooling for sequencing. cDNA library quality and size were measured through a dsDNA High Sensitivity (HS) assay in an Agilent 2100 BioAnalyzer (Agilent Technologies). A quantitative real-time PCR (qPCR) was run to measure cDNA library concentration using the KAPA Library Quantification Kit. qPCR settings were as follows: Initial denaturation (5 min at 95 °C for 1 cycle), then denaturation (30 sec at 95 °C) and annealing/extension/data acquisition (45 sec at 60 °C) combined for 35 cycles. The libraries were then run on the Illumina HiSeq 2500 sequencing platform, with output of paired-end reads of 150 bp by the FAS Center for Systems Biology at Harvard University.

### Sequence processing and de novo assembly

Demultiplexed Illumina HiSeq 2500 sequencing datasets of the 30 tissue samples, in FASTQ format, were retrieved; the quality of the raw reads was assessed and visualized using FASTQC v. 0.11.5 (www.bioinformatics.babraham.ac.uk). Adapter sequences and bases with low-quality phred scores (<30) were trimmed off, and a length filter was applied retaining sequences of >25 bases using TRIMGALORE v. 0.4.2 (www.bioinformatics.babraham.ac.uk).

Two *de novo* transcriptome assemblies for *Syllis magdalena* were constructed with the software Trinity to streamline further differential gene expression analyses (Grabherr et al. 2011; Haas et al. 2013): a reference transcriptome (REFSOM assembly) containing reads from only the somatic parts (anterior part, proventricle, final segments) of each individual of both REPRO and NON-REPRO specimens (23 libraries), and a reference transcriptome including the 5 different parts (anterior part, proventricle, final segments, anterior half part of stolon and posterior half of stolon) of each individual (13 libraries) for only the reproductive specimens (REFTOTREPRO assembly). We did not obtain enough RNA from two of the female tissue samples, proventricle of specimen 0 and anterior part of stolon of specimen 1, to build a library, and therefore conditions ‘proventricle’ and ‘anterior half of stolon’ were represented by a single library in females. Given the large number of raw reads obtained in our study (> 500 million reads), we assembled two different reference transcriptomes, since assembling a single reference transcriptome with the available computational resources would have proved computationally impossible. Raw reads have been deposited in the Sequence Read Archive (BioProject ID PRJNA434571; SRA SUB3709568).

### Transcriptome characterization: Blast and Annotation

Annotation of transcriptome contigs or transcripts (containing all isoforms) for both *de novo* assemblies were done separately using BLASTX against a selection of non-redundant (nr) database from NCBI containing only proteins from Metazoa, with an expected value (E-value) cutoff of 1e^−5^ (Altschul et al.1997). Blast results of the two *de novo* assemblies were used to retrieve Gene Ontology (GO) terms with BLAST2GO 4.0.2 (Conesa *et al.* 2005) under the three different categories: cellular component (CC), biological processes (BP) and molecular function (MF). In addition, GO enrichment analyses using Fisher’s test were done in BLAST2GO, to assess which GO terms were significantly overrepresented in pairwise comparisons between both REFSOM and REFTOTREPRO transcriptomes. The *p*-value for the reciprocal comparisons was adjusted to a 0.05 false discovery rate (FDR) (Benjamini and Hochberg 1995). The Galaxy web-based platform (http://usegalaxy.org) was used to align the RSEM results of each sample with BLASTX results for the *de novo* assemblies for display.

### Estimation of expression levels

In order to obtain expression levels, as read counts, of genes (with all isoforms collapsed) for each tissue type of *S. magdalena* specimens in both reproductive and non-reproductive conditions, trimmed paired reads after trimming were mapped against the reference transcriptome, using BOWTIE2 v. 2.2.1 (Langmead and Salzberg 2012), as implemented in Trinity (Grabherr et al. 2011). The software RSEM v. 1.2.11 (Li and Dewey 2011) was used to generate a table containing read counts.

### Differential gene expression analyses

Differential gene expression analyses were computed in pairwise comparisons of different tissues and conditions using the R package DESeq2, which allows analyses to be performed with low numbers of replicates (Ander and Huber 2010). Before analyzing differential gene expression, read counts were normalized by estimating a scaling factor for each transcript in DESeq2 (Dillies et al. 2013). The significance value for multiple comparisons was FDR adjusted to 0.01 (Benjamini and Hochberg 1995). Visualization of the significant outcomes of genes differentially expressed (upregulated and downregulated) between the tissues and conditions was obtained with a heatmap performed with the ‘GPLOTS’ package of R (http://www.r-project.org/). Using the GO annotation results for the ‘reference’ transcriptome, we obtained the GO terms associated with the differentially expressed isoforms in both pairwise comparisons, which were then implemented together with their *p*-value (adjusted) associated in REVIGO web server (Supek et al. 2011), and graphically represented with the ‘TREEMAP’ function in R. Size of the rectangles was adjusted to reflect the *p*-value using the abs_log_pvalue option in REVIGO.

### Phylogenetic analyses

The evolutionary history of specific genes that could potentially be involved in the stolonization process was also assessed through phylogenetic inference. The translated amino acid sequences of these genes were aligned with orthologues of the same genes in other metazoans obtained from GenBank using MUSCLE ver. 3.6 (Edgar 2004). The G-protein coupled receptors *DAr2* and *SERT* were analyzed together. Both *vasa* and *PL10* are *DEAD-box helicases* and were analyzed together. Other genes were examined in their individual gene families. We selected the best-fit model of amino acid substitution (LG + Γ + G, WAG, as indicated in Figure legends) with ProtTest ver. 2.4 (Abascal et al. 2005) under the Akaike Information Criterion (Posada and Buckley 2004) and later fed into the software for phylogenetic reconstruction. Maximum likelihood analyses of all the genes were conducted in RAxML ver. 7.2.7 (Stamatakis 2006) with 500 independent searches, and 1000 bootstrap replicates (Stamatakis et al. 2008).

## Acknowledgments

The authors are indebted to many members of the Giribet Lab at Harvard University for their help during sample processing, specially to Dr. Sarah Lemer and Dr. David Combosch (currently at University of Guam). Special thanks go to Dr. Greg Rouse (Scripps Institution of Oceanography, UCSan Diego) and Dr. Carlos Sentís (UAM) who provided comments and advice at the beginning of the research. The first author is also very grateful to Dr. Raquel Pérez-Palacios, Dr. Jorge Barbazán (Institut Curie, Paris) and members of Dr. Michel Vervoort lab (Institut Jacques Monod) for their support and useful comments to improve the last version of the manuscript. We also thank also to Milagros Guerra (CBM, CSIC) for his help with TEM observations at UAM. This research received funding from the European Union’s (European Atomic Energy Community’s) Seventh Framework Program (FP7/ 2007–2013; FP7/2007–2011) under grant agreement 227799 to P.A.-C. Sequencing and analyses were conducted with internal MCZ funds to G.G. and with the support of the Center for Systems Biology and the Research Computing group, both from the Faculty of Arts and Sciences, Harvard University.

## Supplementary Material

**Supplementary File S1.** General statistics of the ‘reference’ de novo transcriptomes REFSOM and REFTOTOREP. Total number of trimmed reads used for the assemblies (Total Reads), number of aligned reads for each tissue (Aligned Reads), percentage of aligned reads for each tissue, number of aligned transcripts (Align. Transcr.: including genes and isoforms collapsed into genes), number of genes, percentage og genes, parameter N50, number of transcripts with blast hit against proteins of metazoans (Hit_Mtz), number of annotated transcripts from metazoans (Annot), percentage of GC (GC%), median transcript length (MTL), average transcript length (ATL) and number of assembled bases expressed as Mb.

**Supplementary File S2.** (A) Assigned Gene Ontology (GO) terms for each transcriptome including 3 different levels: CC, BP and MF. (B) GO-term enrichment analysis based on the Fisher’s tests results of pairwise comparisons between both transcriptomes, showing the 36 processes overrepresented in REFSOM, and (C) the 8 processes overrepresented in REFTOTREPRO.

**Supplementary File S3.** (A) Heatmap and (B) correlation matrix of differentially expressed genes from all the tissues and conditions analyzed.

**Supplementary File S4.** Differentially expressed genes with blast hit, and further putative annotations (GO terms), from the pairwise comparisons of somatic tissues (anterior part, proventricle, final segments) between reproductive and non-reproductive individuals.

**Supplementary File S5.** Differentially expressed genes with blast hit, and putative annotations (GO terms), from the pairwise comparisons of somatic tissues (anterior part, proventricle, final segments) and reproductive tissues (stolons) between female and male individuals.

**Supplementary File S6.** Phylogenetic reconstruction of the protein alignment for all the paralogs of the genes *ovochymase* and *chymotripsin* found in our samples. *Ovochymase 1* was differentially expressed in the final segments of reproductive females, and *ovochymase 2* (clearly a *chymotrypsin* in this hypothesis) was differentially expressed in P of reproductive females.

**Supplementary File S7.** Phylogenetic reconstruction of the protein alignment for the DEAD-box helicases *vasa* and *PL10* showing the two homologs of *vasa* (*vasa 1* differentially expressed) and one for *PL10*.

**Supplementary File S8.** Phylogenetic reconstruction of the protein alignment for the different enzymes involved in the synthesis of methyl farnesoate. *Farnesol oxidase/dehydrogenase* (SDR11) (A), *Farnesyl pyrophosphate synthase* (FPP) (B), Farnesyl phosphatase (C), and *Farnesal dehydrogenases* (ALDHE3) (D).

